# Milteforan, a promising veterinary commercial product against feline sporotrichosis

**DOI:** 10.1101/2024.02.14.580352

**Authors:** Laura C. García Carnero, Thaila F. dos Reis, Camila Diehl, Patricia Alves de Castro, Lais Pontes, Camila Figueiredo Pinzan, Gustavo H. Goldman

## Abstract

Sporotrichosis, the cutaneous mycosis most commonly reported in Latin America, is caused by the *Sporothrix* clinical clade species, including *Sporothrix brasiliensis* and *Sporothrix schenckii sensu stricto*. In Brazil, *S. brasiliensis* represents a vital health threat to humans and domestic animals due to its zoonotic transmission. Itraconazole, terbinafine, and amphotericin B are the most used antifungals for treating sporotrichosis. However, many strains of *S. brasiliensis* and *S. schenckii* have shown resistance to these agents, highlighting the importance of finding new therapeutic options. Here, we demonstrate that milteforan, a commercial veterinary product against dog leishmaniasis whose active principle is miltefosine, is a possible therapeutic alternative for the treatment of sporotrichosis, as observed by its fungicidal activity *in vitro* against different strains of *S. brasiliensis* and *S. schenckii*, and by its antifungal activity when used to treat infected epithelial cells and macrophages. Our results suggest milteforan as a possible alternative to treat feline sporotrichosis.

## Introduction

Sporotrichosis, a chronic cutaneous and subcutaneous infection, is the most commonly reported mycosis in Latin America and Asia, with a high prevalence in tropical and subtropical areas, including Brazil, Mexico, Argentina, India, Japan, and China (1, 2). Since 1998, Brazil has experienced large outbreaks of sporotrichosis that have been expanding throughout the country, mainly in the southeastern regions, the reason for which Brazil is considered a hyperendemic area (3–5).

Until 2007, *Sporothrix schenckii* was assumed to be the unique etiological agent for sporotrichosis, but recent molecular analyses have revealed the existence of several cryptic species capable of causing infection (6). These species comprise the *S. schenckii* clinical/pathogenic clade, which includes *S. schenckii sensu stricto*, *S. brasiliensis*, *Sporothrix globosa*, and *Sporothrix lurei* (7, 8). These species are thermodimorphic fungi, with a mycelial phase that grows in decaying organic matter at 25°C (known as the infectious morphology) and a yeast phase that develops inside the host during infection (known as the parasitic morphology) (9, 10). The virulence profile varies among the species of the pathogenic clade being *S. brasiliensis* the most virulent, followed by *S. schenckii,* both with the capacity to cause severe infection even in immunocompetent individuals, while *S. globosa* and *S. lurei* are classified as low virulent species (11, 12).

Sporotrichosis can present different clinical manifestations, such as cutaneous (lymphocutaneous and fixed cutaneous), disseminated cutaneous, and extracutaneous (pulmonary, osteoarticular, ocular, meningeal, and visceral) (13). The development of one or other clinical forms depends on different factors, which include the host immune competence, site and depth of inoculation, amount of inoculum, and the etiological agent, all of which should be considered for proper patient management (14).

The transmission of the *Sporothrix* species is through traumatic implantation with contaminated material, the sapronosis, and the classical route. However, in hyperendemic zones, such as Brazil, zoonotic infection by *S. brasiliensis* is highly reported, transmitted mainly by cats through scratching, biting, and even through contact with fluids from infected animals. This zoonotic transmission is considered a severe health problem in Brazil, especially in the area of Rio de Janeiro, due to the rapid spread of *S. brasiliensis*, which is associated with severe clinical manifestations in both humans and cats (15–18). Besides cats, dogs, albeit to a lesser extent, have also been affected by sporotrichosis, making this infection a significant veterinarian problem. Five thousand hundred-thirteen cases of feline sporotrichosis (from 1988 to 2017) and 244 canine cases (from 1988 to 2014) have been reported by the Evandro Chagas National Institute of Infectious Diseases in Rio de Janeiro, Brazil. However, this number is likely underestimated because sporotrichosis incidence is a mandatory notification only in a few states of Brazil (18).

Identification of the sporotrichosis causative agent is essential for treatment since the *Sporothrix* species show different antifungal susceptibility profiles (19–21), but this is not always possible given that the identification of the species requires molecular tools (8). In general, for the treatment of the cutaneous forms, itraconazole (ITZ) is considered the gold standard for the cutaneous clinical forms, while amphotericin B (AMB) is the first-line antifungal therapy used for disseminated forms (22, 23). However, in the last few years, many *S. brasiliensis* clinical strains have been reported to show resistance to both azoles and AMB (24–26), which complicates sporotrichosis treatment.

Miltefosine (MFS), also known as hexadecyl phosphocholine, is a synthetic glycerol-free phospholipid analog initially used as an antineoplastic drug (27, 28). Nowadays, MFS is the only available oral drug used in the treatment of visceral and cutaneous leishmaniasis in dogs and humans due to its significant antiparasitic activity, *in vitro* and *in vivo*, against *Leishmania* species (29–32). MFS’s action mechanism(s) has yet to be entirely understood. However, it has been demonstrated to act as a multi-target drug associated with the disruption of many vital pathways, such as (i) the inhibition of the biosynthesis of phosphatidylcholine, which causes low levels of this phospholipid (33, 34); (ii) the interference of the cell membrane calcium channels, which induces an increase of intracellular Ca^2+^ (35, 36); (iii) the inhibition of the sphingomyelin biosynthesis, which increases ceramide concentration (37), resulting in cell apoptosis; and (iv) the immune response, in which its immunomodulatory effects induce the activation of the Th1 response, mainly through the increased production of IFNγ and IL-12, which prevails over the Th2 response driven by *Leishmania sp* (38).

MFS has also been reported as an antifungal agent *in vitro* against some of the most clinically significant pathogenic and opportunistic fungi, such as *Candida* spp., *Aspergillus* spp., *Fusarium* spp., and *Cryptococcus* spp. (39–44). In addition, it was recently shown that MFS has *in vitro* fungicidal activity against *Sporothrix* spp., inhibiting the growth of the mycelial phase of *S. brasiliensis*, *S. schenckii*, and *Sporothrix globosa* (45), and the yeast phase of *S. brasiliensis* strains resistant to (ITZ) and AMB (46). It was also demonstrated that alone or in combination with potassium iodide, MFS inhibits the biofilm formation of *S. brasiliensis*, *S. schenckii*, and *S. globosa* (47, 48). All of this evidence suggests the potential of MFS for treating sporotrichosis. Repurposing orphan drugs, which are the application of existing drugs for different therapeutic purposes than the ones initially marketed for, is a good alternative for treating infections caused by susceptible or resistant microorganisms (49). Such is the case of MFS, which, besides being repurposed for treating leishmaniasis, has been recently designated for treating primary amebic meningoencephalitis and invasive candidiasis (50).

Here, we demonstrate that MFS has fungicidal *in vitro* activity against both morphologies (hyphae and yeast) of different *S. brasiliensis* and *S. schenckii* strains. We also showed that milteforan (ML), a commercial veterinary product against dog leshmaniasis whose active principle is miltefosine (Virbac), can inhibit and kill *Sporothrix spp in vitro.* ML treatment also increases the killing of *S. brasiliensis* yeast by the epithelial cells A549 and bone marrow-derived macrophages (BMDMs). Our results suggest ML as a possible veterinary alternative to treat feline sporotrichosis.

## Results

### ML and MFS have fungicidal activity against *Sporothrix* spp. *in vitro*

Several drugs’ *in vitro* antifungal activity against six strains of *S. schenckii* and *S. brasiliensis*, three from each species, were assessed according to their MIC and MFC values for the mycelial and yeast phases (Table 1). From these drugs, ITZ has already been reported to show fungistatic activity against *Sporothrix* spp., while terbinafine (TRB), AMB, and MFS are fungicidal drugs (19, 23, 24). On the other hand, voriconazole (VCZ) was reported to show low activity in inhibiting *Sporothrix* growth, while caspofungin (CSP) does not exhibit antifungal activity *in vitro* (20). We also included brilacidin (BRI), a host defense peptide mimetic that synergizes CSP against several human pathogenic fungi (51), to assess its antifungal activity against *Sporothrix* species.

**Table 1.** MIC and MFC values of several antifungals against *S. schenckii* and *S. brasiliensis* yeast and mycelial phases.

|  |  |  | CSP<br>(4-0.06<br>µg/mL) | VCZ<br>(4-0.06<br>µg/mL) | BRI<br>(80-1.25<br>µM) | ITZ<br>(8-0.125<br>µg/mL) | MFS<br>(16-0.25<br>µg/mL) | ML<br>(16-0.25<br>µg/mL) | TRB<br>(4-0.06<br>µg/mL) | AMB<br>(8-0.125<br>µg/mL) |
| --- | --- | --- | --- | --- | --- | --- | --- | --- | --- | --- |
| Ss<br>4820 | Y | MIC | >4 | >4 | 5 | 0.25 | 2 | 2 | 1 | 2 |
|  |  | MFC | >4 | >4 | 5 | 0.5 | 2 | 2 | 1 | 2 |
|  | M | MIC | >4 | >4 | >80 | 2 | 2 | 2 | 1 | ND |
|  |  | MFC | >4 | >4 | >80 | >8 | 2 | 2 | 1 | ND |
| Ss<br>4821 | Y | MIC | >4 | >4 | 5 | 0.5 | 2 | 2 | 1 | 2 |
|  |  | MFC | >4 | >4 | 5 | 2 | 2 | 2 | 1 | 2 |
|  | M | MIC | >4 | >4 | >80 | 1 | 2 | 2 | 1 | ND |
|  |  | MFC | >4 | >4 | >80 | >8 | 2 | 2 | 1 | ND |
| Ss<br>4822 | Y | MIC | >4 | >4 | 2.5 | 0.125 | 2 | 2 | 0.5 | 2 |
|  |  | MFC | >4 | >4 | 2.5 | 0.25 | 2 | 2 | 0.5 | 2 |
|  | M | MIC | >4 | >4 | >80 | 2 | 2 | 2 | 1 | ND |
|  |  | MFC | >4 | >4 | >80 | >8 | 2 | 2 | 1 | ND |
| Sb<br>4823 | Y | MIC | >4 | >4 | 2.5 | 2 | 2 | 2 | 0.5 | 2 |
|  |  | MFC | >4 | >4 | 2.5 | >8 | 2 | 2 | 0.5 | 2 |
|  | M | MIC | >4 | >4 | >80 | 1 | 2 | 2 | 1 | ND |
|  |  | MFC | >4 | >4 | >80 | >8 | 2 | 2 | 1 | ND |
| Sb<br>4824 | Y | MIC | >4 | >4 | 2.5 | 0.5 | 2 | 2 | 0.5 | >8 |
|  |  | MFC | >4 | >4 | 2.5 | 1 | 2 | 2 | 0.5 | >8 |
|  | M | MIC | >4 | >4 | >80 | 2 | 2 | 2 | 1 | ND |
|  |  | MFC | >4 | >4 | >80 | >8 | 2 | 2 | 1 | ND |
| Sb<br>4858 | Y | MIC | >4 | >4 | 2.5 | 0.125 | 2 | 2 | 0.125 | 2 |
|  |  | MFC | >4 | >4 | 2.5 | 2 | 2 | 2 | 0.125 | 2 |
|  | M | MIC | >4 | >4 | >80 | 1 | 2 | 2 | 1 | ND |
|  |  | MFC | >4 | >4 | >80 | >8 | 2 | 2 | 1 | ND |

Similar to previous reports, we found that none of the *Sporothrix* strains, in either yeast or mycelium states, were inhibited by CSP or VCZ. At the same time, both morphologies from all the isolates were sensitive to low concentrations of TRB and AMB (MIC ≤ 2 µg/mL). For ITZ, all strains’ conidia were highly resistant (MFC > 8 µg/mL). At the same time, the yeast phase was more sensitive with MIC and MFC values ≤ 2 µg/mL, except the *S. brasiliensis* clinical isolate 4823 yeast phase, which shows resistance to the drug (MFC > 8 µg/mL), as already reported (52). In the case of BRI, the yeast morphology from all of the *Sporothrix* strains was susceptible to low concentrations (MIC ≤ 5 µg/mL) of the drug, while conidia are highly resistant. TRB, AMB, and BRI present fungicidal activity against *Sporothrix* species, while ITZ is a fungistatic drug (Table 1). MFS and ML also have fungicidal activity *in vitro* against both morphologies from the *S. schenckii* and *S. brasiliensis* strains, with MIC and MFC values ≤ 2 µg/mL (Table 1 and Figure 1).

**Figure 1.**
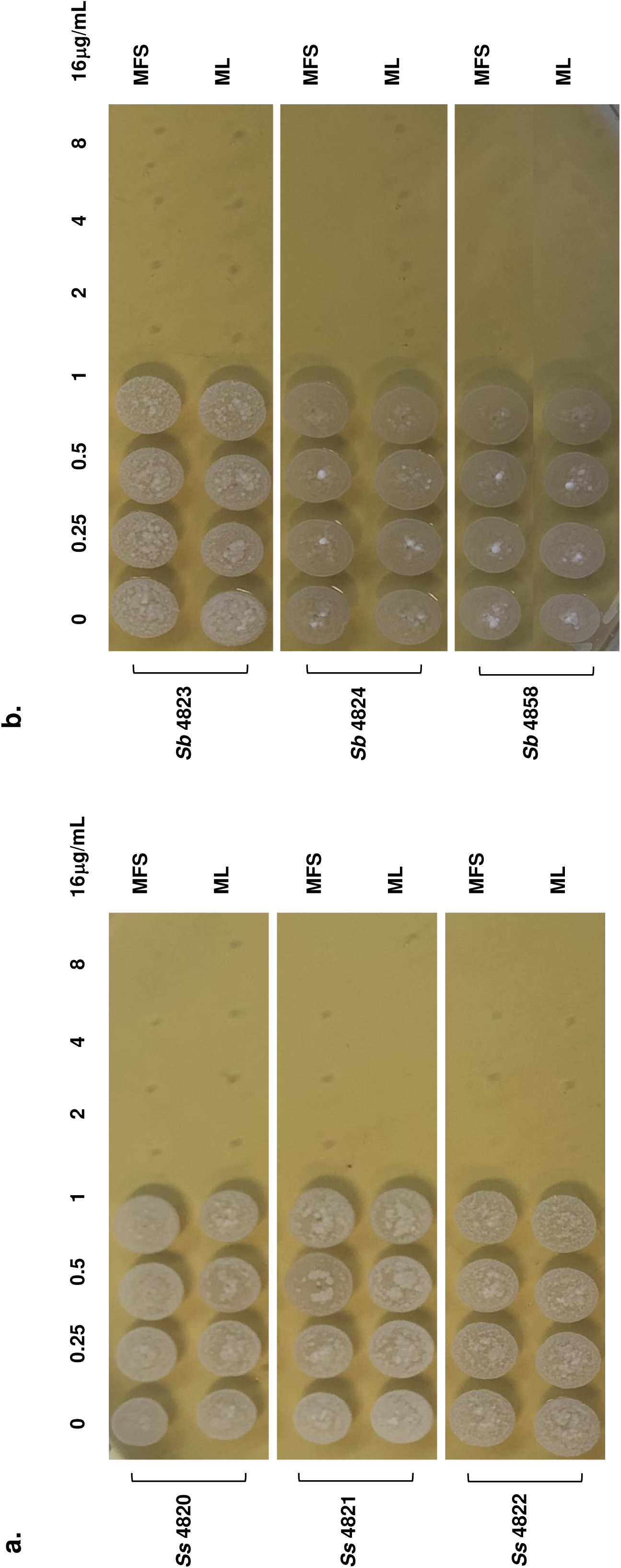
*In vitro* fungicidal activity of miltefosine and milteforan against the yeast morphology of *S. schenckii* and *S. brasiliensis*. a) *S. schenckii* (strains 4820, 4821, and 4822) yeast were grown in liquid YDP pH 7.8 at 37°C in the presence of several concentrations of MFS or ML (16, 8, 4, 2, 1, 0.5, and 0.25μg/mL). After 4 days of incubation, the cells were plated in solid YPD pH 7.8 and incubated for 4 days at 37°C. b) *S. brasiliensis* (strains 4823, 4824, and 4858) yeast were grown in liquid YDP pH 7.8 at 37°C in the presence of several concentrations of MFS or ML (16, 8, 4, 2, 1, 0.5, and 0.25μg/mL). After 4 days of incubation, the cells were plated in solid YPD pH 7.8 and incubated for 4 days at 37°C. As control, yeast cells of each strain were grown without the drugs. Results represent the average of three independent experiments performed by duplicate.

Once we showed the antifungal activity of MFS and ML against *Sporothrix spp.*, we evaluated their ability to interact with some of the drugs already being used for treating sporotrichosis. MIC and MFC values of CSP, VCZ, ITZ, TRB, BRI, and AMB in combination with half MIC of MFS or ML were determined for the yeast morphology of each *Sporothrix* strain (Table 2). No differences in the activity of CSP and VCZ were observed since neither of these drugs could inhibit *S. schenckii* or *S. brasiliensis* growth in the presence of MFS or ML. Combining BRI and MFS or ML does not increase BRI fungicidal activity, as the MIC and MFC values are the same as those of BRI alone. On the other hand, the interaction of MFS or ML with either ITZ, TRB, or AMB increases the antifungal activity against all of the *Sporothrix* strains tested, decreasing their MIC and MFC values.

**Table 2.** MIC and MFC values of MFS and ML combination with several antifungals against *S. schenckii* and *S. brasiliensis* yeast phase.

|  |  |  | CSP<br>(16-0.25µg/mL) | VCZ<br>(16-0.25µg/mL) | ITZ<br>(8-0.125µg/mL) | TRB<br>(4-0.06µg/mL) | BRI<br>(20-0.03µM) | AMB<br>(8µg-0.125µg/mL) |
| --- | --- | --- | --- | --- | --- | --- | --- | --- |
| Ss<br>4820 | Y | MIC | >16 | >16 | 0.25 | 1 | 5 | 2 |
|  |  | MFC | >16 | >16 | 0.5 | 1 | 5 | 2 |
|  | ML | MIC | >16 | >16 | <0.125 | <0.06 | 5 | 1 |
|  |  | MFC | >16 | >16 | 0.5 | <0.06 | 5 | 1 |
|  | MFS | MIC | >16 | >16 | <0.125 | <0.06 | 5 | 1 |
|  |  | MFC | >16 | >16 | 0.5 | <0.06 | 5 | 1 |
| Ss<br>4821 | Y | MIC | >16 | >16 | 0.5 | 0.5 | 5 | 2 |
|  |  | MFC | >16 | >16 | 2 | 0.5 | 5 | 2 |
|  | ML | MIC | >16 | >16 | <0.125 | 0.25 | 5 | 0.5 |
|  |  | MFC | >16 | >16 | 0.5 | 0.25 | 5 | 0.5 |
|  | MFS | MIC | >16 | >16 | <0.125 | 0.25 | 5 | 0.5 |
|  |  | MFC | >16 | >16 | 0.5 | 0.25 | 5 | 0.5 |
| Ss<br>4822 | Y | MIC | >16 | >16 | 0.125 | 0.5 | 5 | 2 |
|  |  | MFC | >16 | >16 | 0.25 | 0.5 | 5 | 2 |
|  | ML | MIC | >16 | >16 | ND | ND | 5 | 1 |
|  |  | MFC | >16 | >16 | ND | ND | 5 | 1 |
|  | MFS | MIC | >16 | >16 | ND | ND | 5 | 1 |
|  |  | MFC | >16 | >16 | ND | ND | 5 | 1 |
| Sb<br>4823 | Y | MIC | >16 | >16 | 2 | 0.5 | 2.5 | 2 |
|  |  | MFC | >16 | >16 | >8 | 0.5 | 2.5 | 2 |
|  | ML | MIC | >16 | >16 | 0.5 | 0.125 | 2.5 | 0.5 |
|  |  | MFC | >16 | >16 | >8 | 0.125 | 2.5 | 0.5 |
|  | MFS | MIC | >16 | >16 | 0.5 | 0.125 | 2.5 | 0.5 |
|  |  | MFC | >16 | >16 | >8 | 0.125 | 2.5 | 0.5 |
| Sb<br>4824 | Y | MIC | >16 | >16 | 0.5 | 0.5 | 2.5 | >8 |
|  |  | MFC | >16 | >16 | 1 | 0.5 | 2.5 | >8 |
|  | ML | MIC | >16 | >16 | 0.25 | 0.25 | 2.5 | 8 |
|  |  | MFC | >16 | >16 | 0.25 | 0.25 | 2.5 | 8 |
|  | MFS | MIC | >16 | >16 | 0.25 | 0.25 | 2.5 | 8 |
|  |  | MFC | >16 | >16 | 0.25 | 0.25 | 2.5 | 8 |
| Sb<br>4858 | Y | MIC | >16 | >16 | 0.125 | 0.125 | 5 | 2 |
|  |  | MFC | >16 | >16 | 2 | 0.125 | 5 | 2 |
|  | ML | MIC | >16 | >16 | ND | ND | 5 | 0.25 |
|  |  | MFC | >16 | >16 | ND | ND | 5 | 0.25 |
|  | MFS | MIC | >16 | >16 | ND | ND | 5 | 0.25 |
|  |  | MFC | >16 | >16 | ND | ND | 5 | 0.25 |

Ss: *S. schenckii*, Sb: *S. brasiliensis*; Y: untreated yeasts, ML: yeast treated with milteforan (1µg/mL), MFS: yeast treated with miltefosine (1µg/mL); CSP: caspofungin, VCZ: voriconazole, ITZ: itraconazole, TRB: terbinafine, BRI: brilacidin, AMB: amphotericin B; ND: Not determined.

Next, in order to determine what kind of interaction MFS has with ITZ, TRB, and AMB, the drug combination responses were analyzed using checkerboard assays and the SynergyFinder software (53), which evaluates the potential synergy of 2 or more drugs. The dose-response data obtained for combining MFS with either TRB, ITZ, or AMB against *S. brasiliensis* and *S. schenckii* yeast cells shows a likely additive interaction (synergy score from −10 to 10) (Figure 2). As previously reported for ITZ (46), we found that MFS does not synergize with the drug against *S. brasiliensis* and *S. schenckii*.

**Figure 2.**
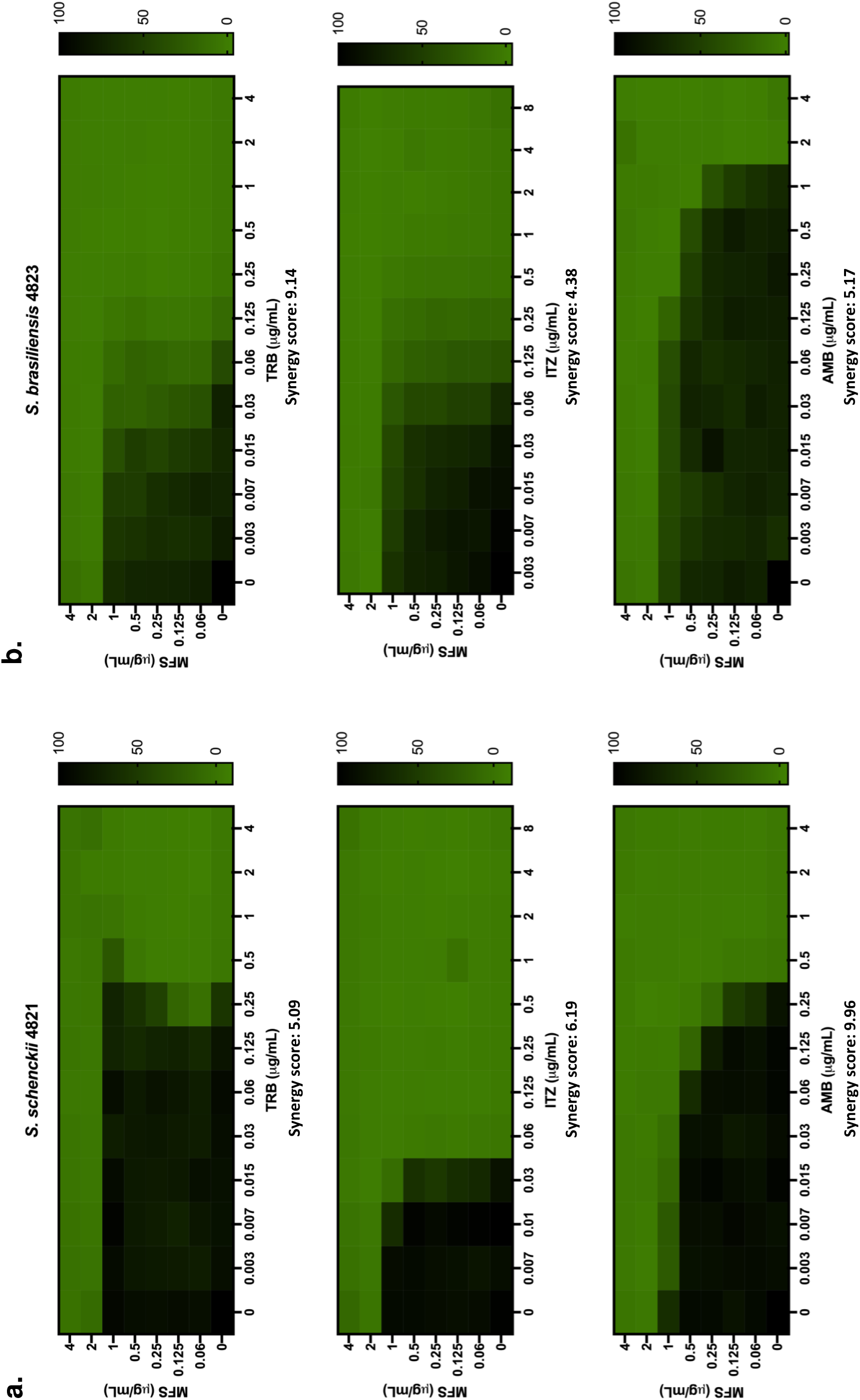
MFS has an additive interaction with ITZ, TRB, and AMB against *S. brasiliensis* and *S. schenckii* yeast cells. The synergy score for MFS x TRB, MFS x ITZ, and MFS x AMB against *Sporothrix* was determined by analyzing the SynergyFinder software’s checkerboard data. a) *S. schenckii* and b) *S. brasiliensis* yeast were grown in liquid YDP pH 7.8 at 37°C in different concentrations of the selected drugs. After 4 days of incubation, the metabolic activity of the cells was assessed by the XTT reduction assay. Results are expressed as the % of metabolic activity and represent the average of three independent experiments.

### MFS localizes to the *Sporothrix* cell membrane and mitochondria and causes cell death

Although the antifungal effect of MFS against *Sporothrix* has been reported, the localization of the drug in the yeast is still unknown. In *Leishmania* (54) and *A. fumigatus* (43), MFS localizes in the cell membrane and the mitochondria, increasing mitochondrial fragmentation and damage. Here, we found that in *S. brasiliensis*, fluorescent MFS is also localized in the cell membrane and the mitochondria in 47% of the cells investigated (three repetitions of 100 cells each), as shown by MitoTracker colocalization (Figure 3).

**Figure 3.**
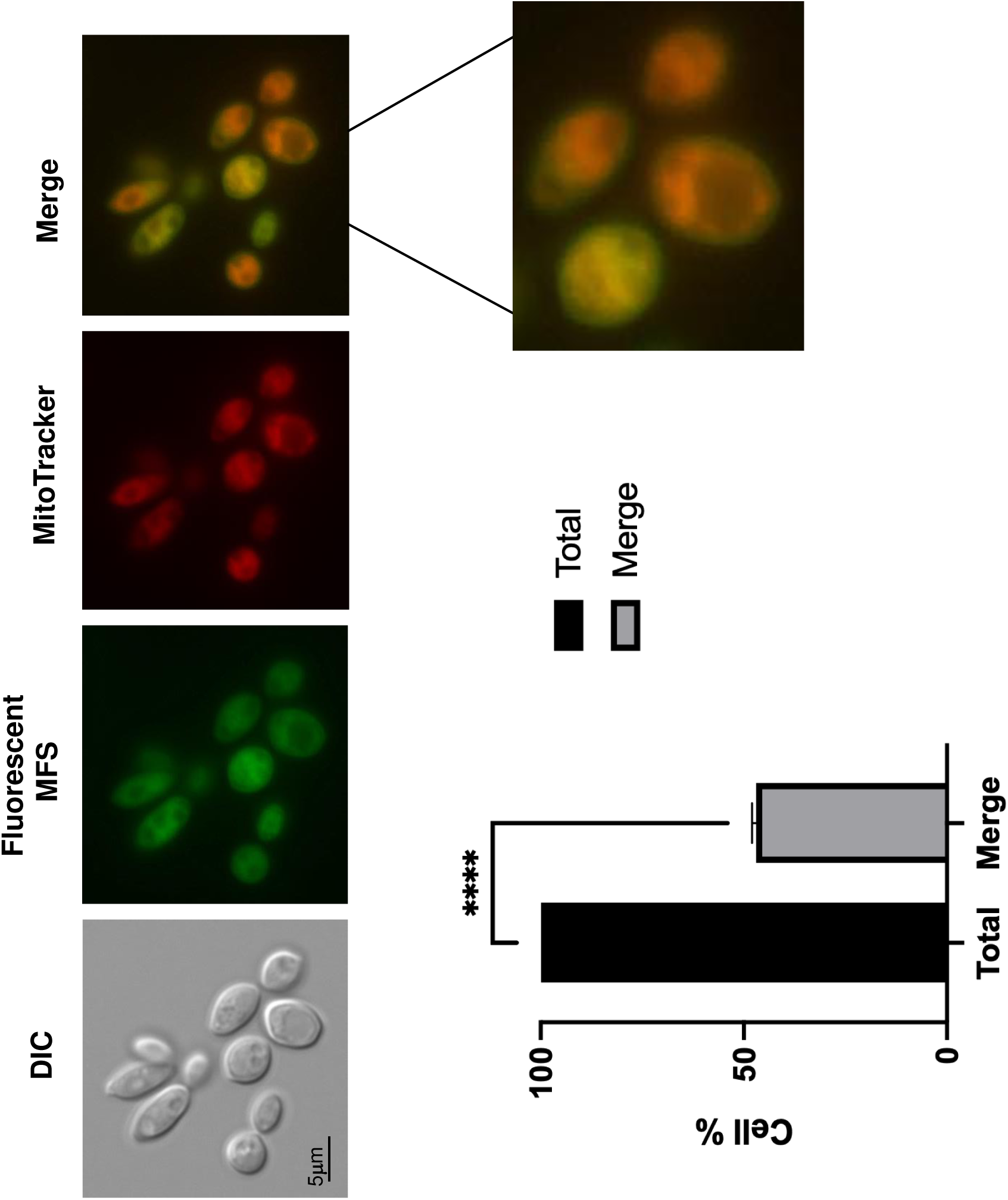
MFS is localized in the mitochondria and cell surface of *S. brasiliensis* yeast. *S. brasiliensis* yeast cells were exposed to fluorescent MFS (2μg/mL) for 1 h and then stained with MitoTracker Deep Red FM. The MFS and MitoTracker signals merge on the mitochondria, while the MFS signal is observed on the cell surface. Three independent experiments were performed, and 100 cells were counted for each to calculate a 47.06 ± 1.01 % of MFS and MitoTracker colocalization (merge).

Subsequently, to evaluate the viability of the yeast in the presence of MFS, drug-treated cells were stained with propidium iodide (PI) and analyzed by fluorescence microscopy. Since PI only penetrates cells with damaged membranes, PI^+^ cells are considered to be going through late apoptosis or early necrosis (55). Treatment of *S. brasiliensis* yeasts with 2, 4, and 8μg/mL of MFS showss dose-dependent damage of the cells since the PI signal increased with the drug concentration (Figure 4), as early as 6 hours of exposure, confirming the MFS fungicidal activity against *Sporothrix*.

**Figure 4.**
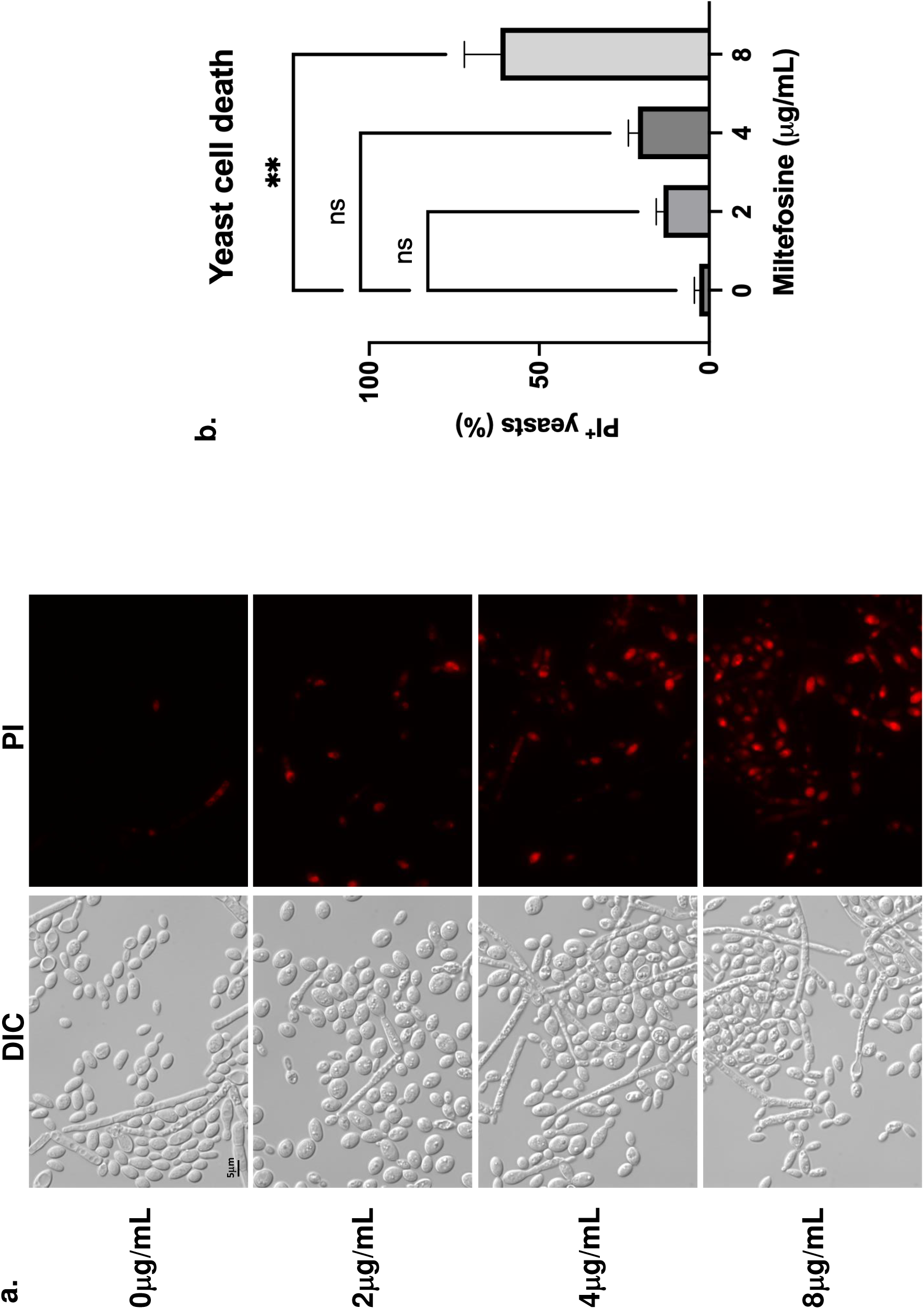
MFS causes dose-dependent death in *S. brasiliensis* yeast. a) *S. brasiliensis* yeast were exposed to 0, 2, 4, and 8μg/mL of MFS for 6 hours, stained with PI, and analyzed by fluorescence microscopy. b) Quantification of PI^+^ yeast exposed to MFS, in which 100 yeast-like cells were counted for each condition. Results represent the average of two independent experiments. \*\**p*-value<0.001 when compared to untreated cells. ns: not significant.

### ML decreases *S. brasiliensis* fungal burden in A549 pulmonary cells and bone marrow-derived macrophages (BMDM)

To determine the antifungal activity of ML against *S. brasiliensis* in the host tissues, two cell lines were used: lung A549 cells and Bone Marrow-Derived Macrophages (BMDMs). As shown in Figure 5a, ML concentrations of 40μg/mL and lower did not reduce A549 cell viability compared to the control. A549 cells were challenged with 1:10 and 1:20 ratios (A549-yeast), and we observed a significant reduction of more than 90 % in the fungal viability in both ML treatments, which contrasts with TRB treatment that shows about 50 % viability (Figure 5b).

**Figure 5.**
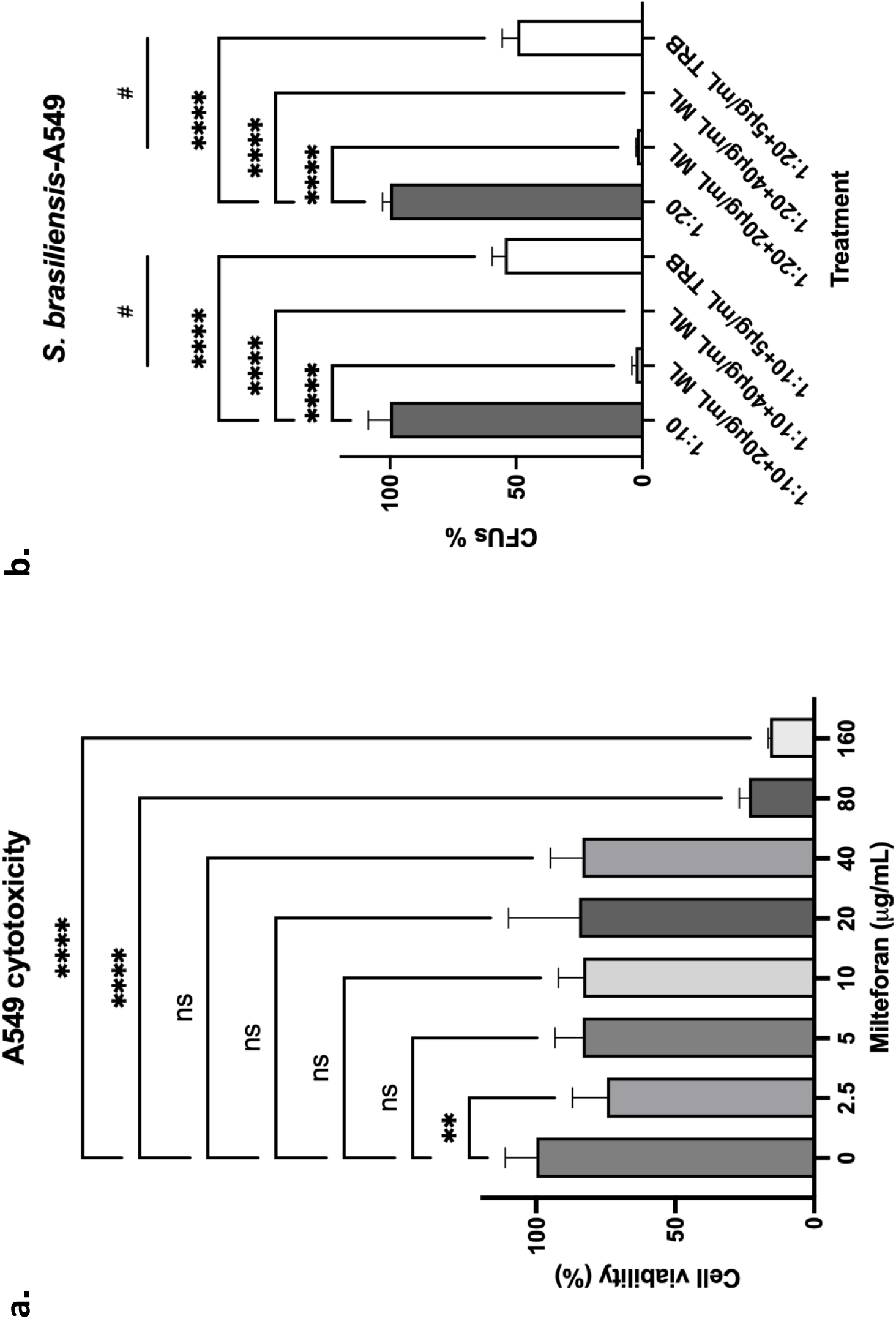
Concentrations up to 40μg/mL of ML are not toxic to human cells and can significantly decrease *S. brasiliensis* survival in A549 epithelial cells. a) A459 epithelial cells were treated with different ML concentrations, with a decrease of cell viability only at 80μg/mL or higher concentrations. b) A459 cells were challenged with *S. brasiliensis* yeast at a proportion of 1:10 and 1:20 and then treated with 20 and 40μg/mL of ML. The fungicidal drug TRB was included as a control. \*\**p*-value<0.01 when compared to untreated cells. \*\*\*\**p*-value<0.0001 when compared to untreated cells. #*p*-value<0.0001 when compared to cells treated with TRB.

When we challenged BMDMs with *S. brasiliensis* at a 1:10 ratio (BMDMs-yeast) in the presence of 20 and 40μg/ml ML, we observed complete clearing of *S. brasiliensis* compared to TRB that showed about 80 and 40 % clearing, respectively, at 24 and 48 h (Figure 6). Our results strongly indicated that ML can help both A549 and BMDMs to clear *S. brasiliensis* infection.

**Figure 6.**
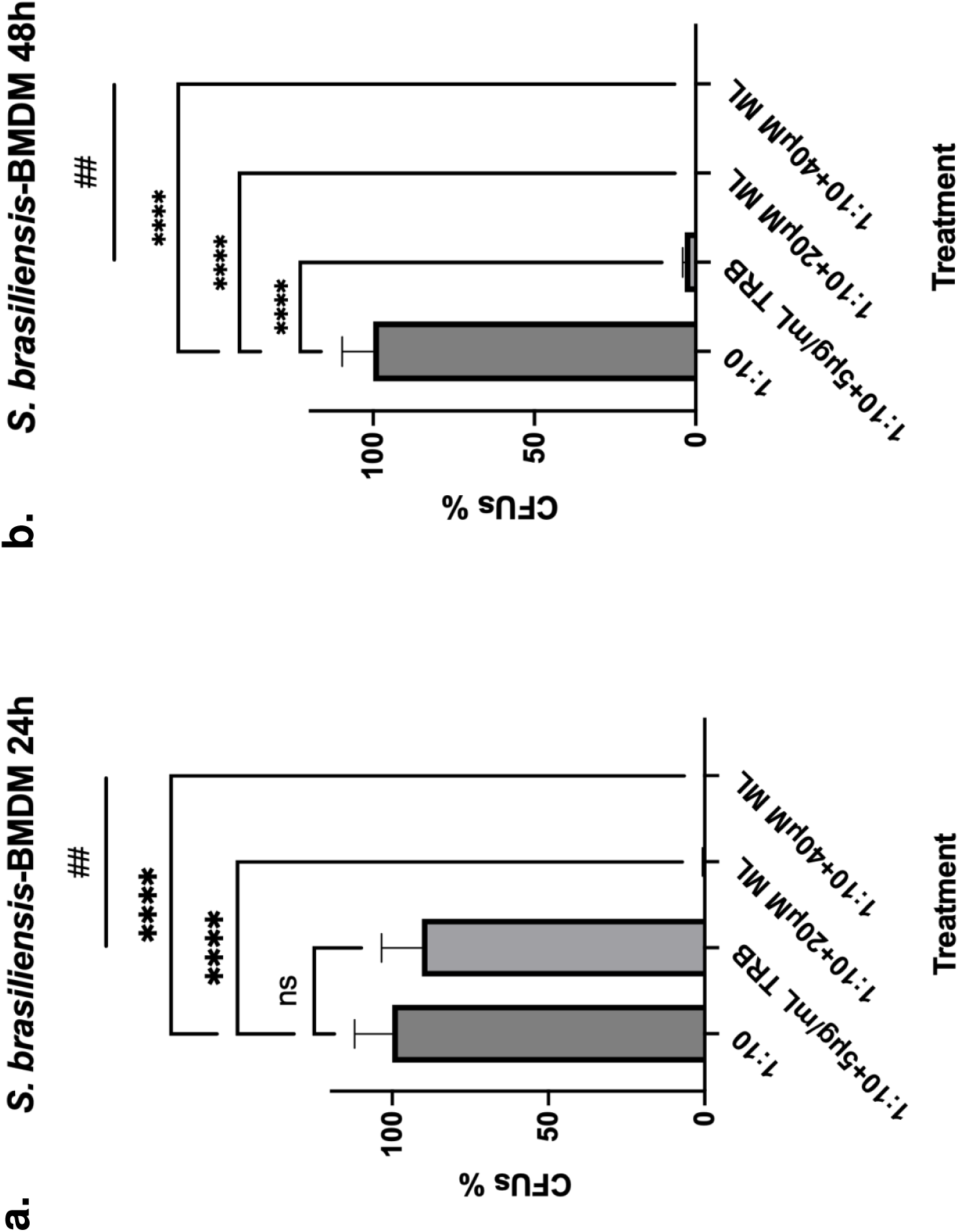
Killing of *S. brasiliensis* yeast by BMDM is significantly increased in the presence of ML. a) BMDM cells were infected with *S. brasiliensis* yeast and then treated with 20 and 40μg/mL for 24h, which decreased the fungal survival by almost 100% compared to untreated cells. b) BMDM cells were infected with *S. brasiliensis* yeast and were then treated with 20 and 40μg/mL for 48h, which decreased the fungal survival to 100% when compared to untreated cells. The fungicidal drug TRB was included as a control. \**p*-value<0.05 when compared to untreated cells. \*\*\**p*-value<0.0005 when compared to untreated cells. \*\*\*\**p*-value<0.0001 when compared to untreated cells. #*p*-value<0.01 when compared to cells treated with TRB. ##*p*-value<0.01.

We also assessed the ability of the BMDMs to produce cytokines after stimulation by *S. brasiliensis* and treatment with the drug. It has already been reported that *S. brasiliensis* yeast stimulates higher production of TNF-α, IL-6, IL-1β, and IL-10 in human monocyte-derived macrophages when compared to *S. schenckii*, and it is also more phagocytosed under these conditions (56), which might contribute to the higher virulence of this species.

After infection of BMDMs and treatment during 24h, we observed a significant decrease in the stimulation of TNF-α and IL-6 when the yeast cells were treated with TRB and 20 and 40μg/mL of ML, when compared to untreated cells (1:10) (Figure 7a). However, when compared to TRB treatment, a significant decrease was observed in the stimulation of TNF-α only at 40μg/mL of ML. In contrast, no difference was observed in the case of IL-6 with both ML concentrations compared to TRB. Finally, for the secretion of IL-10, a significant decrease was only observed when the yeast cells were treated with both ML concentrations. However, no difference was found with the TRB treatment compared to untreated cells (Figure 7a).

**Figure 7.**
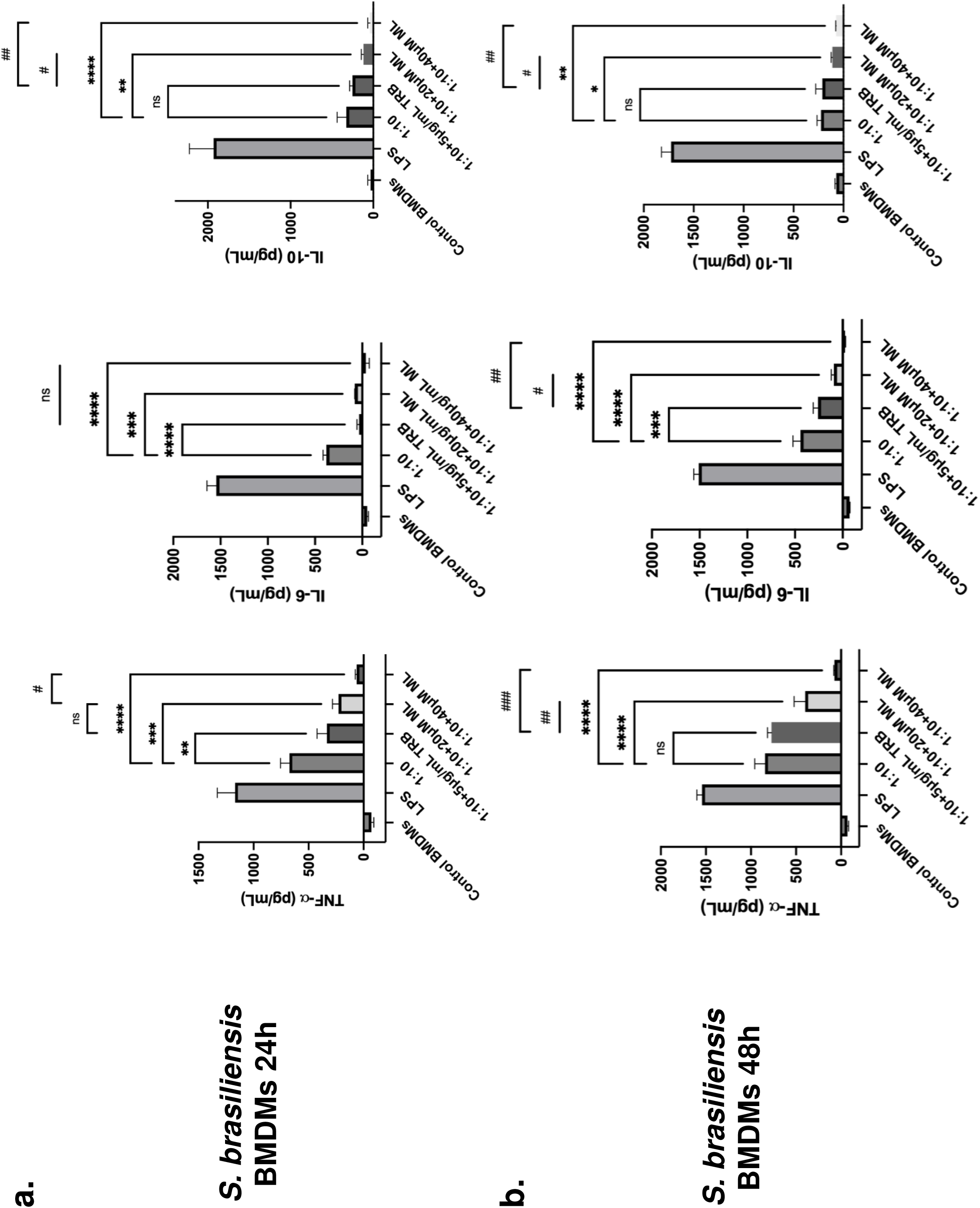
Cytokine secretion by BMDM infected with *S. brasiliensis* and treated with ML. a) BMDM cells were infected with *S. brasiliensis* yeast and treated with ML 20 and 40μg/mL for 24h. The interaction supernatant was collected and the cytokines TNF-α (ns: not significant; \*\**p*-value<0.005 when compared to untreated cells; \*\*\**p*-value<0.0005 when compared to untreated cells; \*\*\*\**p*-value<0.0001 when compared to untreated cells; #*p*-value<0.01 when compared to TRB treatment), IL-6 (ns: not significant; \*\*\**p*-value<0.0005 when compared to untreated cells; \*\*\*\**p*-value<0.0001 when compared to untreated cells), and IL-10 (ns: not significant; \*\**p*-value<0.005 when compared to untreated cells; \*\*\*\**p*-value<0.0001 when compared to untreated cells; #*p*-value<0.0005 when compared to TRB treatment; ##*p*-value<0.0001 when compared to TRB treatment) were measured. b) BMDM cells were infected with *S. brasiliensis* yeast and treated with 20 and 40μg/mL for 48h. The interaction supernatant was collected and the cytokines TNF-α (ns: not significant; \*\*\*\**p*-value<0.0001 when compared to untreated cells; #*p*-value<0.0005 when compared to TRB treatment; ##*p*-value<0.0001 when compared to TRB treatment), IL-6 (\*\*\**p*-value<0.001 when compared to untreated cells; (\*\*\*\**p*-value<0.0001 when compared to untreated cells; #*p*-value<0.005 when compared to TRB treatment; ##*p*-value<0.0001 when compared to TRB treatment), and IL-10 (ns: not significant; \**p*-value<0.05 when compared to untreated cells; \*\**p*-value<0.01 when compared to untreated cells; #*p*-value<0.05 when compared to TRB treatment; ##*p*-value<0.01 when compared to TRB treatment) were measured.

After 48h of infection, treatment with TRB did not cause a significant decrease in the TNF-α production, while both ML concentrations did when compared to untreated cells and TRB treatment (Figure 7b). In the case of the IL-6 secretion, the same trend as that of 24h was observed, with the only exception that treatment with 20 and 40μg/mL of ML results in a significant decrease compared to TRB (Figure 7b). The secretion of IL-10 did not decrease with the TRB treatment, while significantly decreased in macrophages infected and uninfected treated with ML, confirming the participation of this drug in the immune response modulation (Figure 7b).

## Discussion

Although there are several therapeutic options for the treatment of sporotrichosis, fungal resistance and cytotoxicity of the drugs to the host are essential obstacles that hinder the efficient recovery of the patient. ITZ is considered the first-line treatment, an azole known for its fungistatic activity against *Sporothrix* species (22, 24), which has increased the development of resistance in some isolates, mainly from *S. brasiliensis* (46, 57, 58). TRB, a drug with fungicidal activity against *Sporothrix,* has been reported to be effective in treating the cutaneous forms but not for the disseminated infections for which AMB is used. AMB is considered a second-line treatment and is commonly used to treat the invasive and disseminated forms, with the disadvantage that it is very toxic in the doses and time needed to eradicate the infection, in addition to recent reports of isolates resistant to this antifungal agent (22, 46).

In Brazil, cat-transmitted sporotrichosis, caused by *S. brasiliensis*, is a vital health treat that has been spreading since 1998 (5, 8) across the country, affecting domestic animals and humans, another reason for which is of great importance to find new drugs for the treatment and control of this mycosis. For this objective, drug repurposing is an excellent alternative to finding new treatments since these drugs already approved to be used in humans and animals, initially developed to treat other diseases, can help treat infections caused by different pathogens (59, 60). Such is the case of commercial MFS, which was initially used as an antineoplastic drug (27, 28) that is now the only available oral treatment for leishmaniasis in dogs and humans (29–32), and was recently proven to be effective for the treatment of infections caused by *Candida* species (39, 40). As previously demonstrated (45, 46, 48), MFS also has *in vitro* fungicidal activity against *Sporothrix* species by inhibiting the growth of both fungal morphologies. *S. brasiliensis* and *S. schenckii* strains are sensitive to low concentrations of this drug, with an antifungal activity of 2μg/mL for both hyphae and yeast cells. Unlike ITZ, we found no strain resistant to MFS or ML.

We also assessed the ability of MFS to synergize with other drugs used for the treatment of sporotrichosis, including TRB, ITZ, and AMB, and as previously reported for ITZ (45), MFS does not synergize the activity of other antifungals. However, it has instead an additive effect, which suggest they do not interact, or act on independent pathways (61). Similarly to *A. fumigatus* (43), MFS is directed to the mitochondria of *S. brasiliensis* yeast, staying also on the cell surface and causing cell death, suggesting that this drug might be affecting the mitochondria and membrane integrity, which might be related to its mechanism of action.

This drug has been reported to be toxic in high doses in mice, with high mortality in concentrations higher than 25mg/kg (62, 63), with maximum concentrations in the kidney and liver, probably due to its amphiphilic nature (64, 65). We assessed ML cytotoxicity in A549 human pulmonary cells and observed a significant viability reduction at 80μg/mL. When we tested the ability of ML to decrease the fungal burden in A549 cells and BMDMs, at 24h and 24 and 48h, respectively, we observed that ML could significantly decrease the CFUs more efficiently than the fungicidal drug TRB in both cell types, with an almost complete clearing of the yeast cells as early as 24 h of treatment.

One of the proposed mechanisms of action for MFS is its immunomodulatory ability, which is essential for the treatment of leishmaniasis since the drug induces the Th1 response and suppresses the Th2, by increasing the production of proinflammatory cytokines such as IFNγ, TNFα, and IL-12 for the clearance of intracellular pathogens, while relapses of leishmaniasis have been related with an increase of the Th2 response and the production of IL-10 (32, 38). We observed that ML decreases the fungal burden and the production of TNFα, IL-6, and IL-10, secreted by the infected BMDMs. We propose three non-excluded hypotheses to explain it: (i) The cytokines reduction might be related to the fact that the drug is killing the yeast cells before being phagocytosed, where there is the death of the yeast cells as early as 6 hours of MFS treatment; (ii) since the drug is localized to the cell surface, MFS could act as an opsonizing agent helping in the macrophage recognition and further phagocytosis: and (iii) MFS could bind to essential virulence factors, such as adhesins, or immunogenic components, such a β-glucans, in a way that is attenuating *S. brasiliensis* ability to infect and generate an immune response. All three options would reduce the fungal load, tissue damage, and inflammation, making this veterinary drug a suitable treatment alternative for feline sporotrichosis.

## Materials and Methods

### Fungal strains and culture conditions

In this study, three *Sporothrix schenckii* (ATCC-MYA 4820, ATCC-MYA 4821, and ATCC-MYA 4822) and three *S. brasiliensis* strains (ATCC-MYA 4823, ATCC-MYA 4824, and ATCC-MYA 4858) were used for the *in vitro* antifungal susceptibility assays; *S. schenckii* ATCC-MYA 4821 and *S. brasiliensis* ATCC-MYA 4823 were used for the checkerboard assays; and *S. brasiliensis* ATCC-MYA 4823, a highly virulent clinical isolate obtained from feline sporotrichosis (66), was used for the infection assays.

The mycelial phase from *Sporothrix* spp. was obtained and maintained on solid YPD pH 4.5 (yeast extract 1% (w/v), gelatin peptone 2% (w/v), and dextrose 3% (w/v)) at 28°C for four days. In contrast, the yeast morphology was grown in liquid YPD pH 7.8, at 37°C under orbital agitation for four days, as previously reported (67). Each phase was confirmed by observing the cells with light microscopy.

### Antifungal drugs

For the *in vitro* assays, voriconazole (VCZ, Sigma-Aldrich), itraconazole (ITZ, Sigma-Aldrich), amphotericin B (AMB, Sigma-Aldrich), terbinafine (TRB, Sigma-Aldrich), and brilacidin (BRI, supplied by Innovation Pharmaceuticals) were diluted in dimethyl sulfoxide (DMOS); while miltefosine (MFS, Sigma-Aldrich), the milteforan active compound, was diluted in ethanol; and caspofungin (CSP, Sigma-Aldrich) was diluted in distilled water. Milteforan (miltefosine 2%) was purchased from Virbac as an oral solution.

### *In vitro* antifungal susceptibility testing

The minimum inhibitory concentrations (MICs) were determined by the broth microdilution method adapted from protocols published by the Clinical Laboratory Standard Institute for the mycelial and yeast phases (24, 68). Briefly, serial two-fold dilutions of the antifungal drugs were performed in YPD pH 4.5 and 7.8, for mycelial and yeast, respectively, into 96-well microtiter plates to obtain concentrations of 4-0.06μg/mL for CSP, VCZ and TRB; 8-0.125μg/mL for ITZ and AMB; 16-0.25μg/mL for MFS and ML; and 80-1.25μM for BRI, with a final concentration of 2×10^3^ and 2×10^4^ conidia or yeast cells, respectively, in a volume of 100μL. The plates were incubated at 28°C (for conidia) or 37°C (for yeast) for four days, and the MIC was determined by visual inspection and defined as the lowest concentration that inhibits 90-100% of fungal growth about untreated cells. Finally, 5μL of conidia or yeast cells from each well were grown in drug-free solid YPD pH 4.5 and pH 7.8 at 28°C and 37°C, respectively, for four days. The minimum fungicidal concentration (MFC) value was the lowest concentration, showing no fungal growth. Three independent experiments were performed by duplicate.

### Checkerboard assays and synergy testing

The drug combination effect was determined through the MIC and MFC values of the yeast phase, as described before. Briefly, serial twofold dilutions of the antifungal drugs were performed in liquid YPD pH 7.8 containing half MIC of MFS or ML (1μg/mL) in 96-well microtiter plates to obtain concentrations of 16-0.25μg/mL for CSP and VCZ; 8-0.125μg/mL for ITZ and AMB; 4-0.06μg/mL for TRB; and 80-1.25μM for BRI, with a final concentration of 2×10^4^ yeast, in a volume of 100μL. The plates were incubated at 37°C for four days, and the MIC was determined by visual inspection. It was defined as the lowest concentration inhibiting 90-100% of fungal growth in cells treated only with 1μg/mL of MFS or ML. After MIC determination, 5μL of yeast from each well were grown in drug-free solid YPD pH 7.8 at 37°C for four days. The MFC value was the lowest concentration, which showed no fungal growth.

Checkerboard assays were performed to quantify the interaction (synergistic, additive, or antagonistic) between MFS and ITZ, AMB, or TRB. Briefly, a stock solution of 2×10^5^ yeast/mL and each drug (8μg of MFS and 16μg/mL of ITZ, 16μg of AMB, or 8μg of TRB) were prepared in RMPI-1640. In 96-well microtiter plates, the first antibiotic (MFS) was diluted sequentially along the ordinate. In contrast, the second drug (ITZ, AMB, or TRB) was diluted along the abscissa to obtain a final volume of 100μL. The plates were incubated at 37°C for four days, and the metabolic activity was determined through the XTT reduction assay (47). Briefly, 50μL of a solution of XTT 1mg/mL and menadione 1mM resuspended in water were added to each well, mixed, and incubated in the dark at 37°C for three h. The supernatant of each well was transferred to a new plate and read in a spectrophotometer at 492nm. Results are expressed as means ± SD of three independent experiments.

To determine the type of drug interaction, the SynergyFinder software (53) was used, with the following parameters: detect outliners: yes; curve fitting: LL4; method: Bliss; correction: on. The summary synergy scores represent the average excess response due to drug interaction, in which a value less than −10 suggest an antagonistic interaction between two drugs; values from −10 to 10 suggest an additive interaction; and values larger that 10 suggest a synergistic interaction.

### Yeast cells death

The effect of ML on the cell membrane potential was assessed by staining with propidium iodide (PI). Yeast cells grown for 4 days in liquid YPD pH 7.8 were treated with 0, 2, 4 and 8μg/mL of ML during 6 h, stained with PI 20mM for 30 minutes, and washed with PBS 1X three times. Fluorescence was analyzed at an excitation wavelength of 572/25nm and emission of 629/62nm with the Observer Z1 fluorescence microscope using a 100x oil immersion lens objective. Differential interference contrast (DIC) and fluorescent images were capture with an AxioCam camera (Carl Zeiss) and processed using AxioVision software (version 4.8). The experiment was performed twice, and for each treatment at least 100 cells were counted. The results were plotted using Graphpad Prism (GraphPad software, Inc.). A *p*-value<0.001 was considered significant.

### Miltefosine localization

*S. brasiliensis* yeast cells cultured for 4 days in YPD pH 7.8 were washed 3 times with PBS 1X and then treated with the fluorescent MFS analogue MT-11 C-BDP (excitation wavelength 450-490nm and emission wavelength 500-550nm) for 6 hours, also in liquid YPD pH 7.8. The cells were washed 3 times and stained with 250nM of MitoTracker Deep Red FM (Invitrogen) (wavelength absorbance/emission 644/665nm) and washed again. The yeast cells were visualized in slides with the Observer Z1 fluorescent microscope using a 100x oil immersion lens objective. DIC and fluorescent images were capture with an AxioCam camera (Carl Ziess) and processed using AxioVision software (version 4.8). Two independent experiments were performed, and 100 cells were counted of each to calculate the merge %.

### Cytotoxicity assay

The cytotoxicity of ML was determined in A549 human lung cancer cells using the XTT reduction assay. 2×10^5^ cells/well were seeded in 96-well tissue plates and incubated in Dulbecco’s Modified Eagle Medium (DMEM, ThermoFischer). After 24 h of incubation with CO_2_ 5%, the cells were treated with different concentrations of ML (0, 2.5, 5, 10, 20, 40, 80 and 160μg/mL), and after 48 h of incubation, cell viability was assessed using the XTT assay. Briefly, 80μL of a solution of XTT 1mg/mL in DMEM, HEPES 1M, and menadione 8μg/mL were added to each well, and after 30 min, formazan formation was quantified spectrophotometrically at 450nm using a microplate reader. Each treatment was performed by triplicate and the results were plotted using Graphpad Prism (GraphPad Software, Inc.). A *p*-value<0.0001 was considered significant.

### A549 and bone marrow derived macrophages (BMDMs) killing assays

The cell line A549 and BMDMs were cultured using DMEM supplemented with fetal bovine serum (FBS) 10% and penicillin-streptomycin 1% (Sigma-Aldrich), and seeded at a concentration of 1×10^6^ cells/mL in 24-well plates (Corning). The cells were challenged with *S. brasiliensis* yeasts at a multiplicity of infection of 1:10 and were then treated with ML 20 and 40μM. As control, we included untreated cells and cells treated with TRB 5μg/mL. For the BMDMs, cells treated with LPS were also included as control. The A549 were incubated during 24 h at 37°C with CO_2_ 5%, while the BMDM were incubated for 24 and 48h under the same conditions. After incubation, the culture media was removed, each well was washed 3 times with PBS 1X, and 1mL of sterile cold water was added to recover and collect the cell monolayer. To assess the number of CFUs, 100μL of the cell suspensions were plated on YDP pH 4.5 and incubated at 28°C for 4 days. When necessary, the cell suspensions were diluted at 1:100 or 1:1000 and 100μL were plated. 50μL of the inoculum adjusted to 1×10^3^ cells/mL was also plated to correct the CFU count. Each treatment was performed by triplicate to calculate the CFU %, and the results were plotted using Graphpad Prism (GraphPad Software, Inc.). A *p*-value<0.0001 was considered significant.

### Cytokines quantification

The Elisa-assay kits (R&D Systems) were used to evaluate the concentration of the proinflammatory cytokines TNFα and IL-6, and the anti-inflammatory cytokine IL-10 in the supernatants of the *S. brasiliensis* and BMDMs interaction for 24 and

48 h, according to the manufacturers instruction. The plates absorbance was read at 450nm and the cytokine concentration (pg/mL) was calculated according to the values obtained in the standard curve of each cytokine. The results were plotted using Graphpad Prism (GraphPad software, Inc.).

### Statistical analyses

The GraphPad Prism 10 (GraphPad Software, Inc.) was used for the statistical analyses. The results are reported as the media ± SD from two or three independent experiments performed by duplicate and were analyzed using the Ordinary one-way ANOVA or the Unpaired T test. The statistical significance was considered with a *p*-value<0.05 or lower.

## Acknowledgements

We thank the Fundação de Amparo à Pesquisa do Estado de São Paulo (FAPESP) grant numbers 2021/04977-5 (GHG) and 2022/08556-7 (LCGC) 2022/08796-8 (CD), 2022/09882-5 (LP), the Conselho Nacional de Desenvolvimento Científico e Tecnológico (CNPq), FAPESP and Fundação Coordenação de Aperfeiçoamento do Pessoal do Ensino Superior (CAPES) grant number 405934/2022-0 (The National Institute of Science and Technology INCT Funvir), and CNPq 301058/2019-9 from Brazil to GHG., both from Brazil, the National Institutes of Health/National Institute of Allergy and Infectious Diseases from the USA, grant R01AI153356 to GHG. This work was also funded by the Joint Canada-Israel Health Research Program, jointly supported by the Azrieli Foundation, Canada’s International Development Research Centre, Canadian Institutes of Health Research, and the Israel Science Foundation (GHG).

